# Effects of a Cancer-Associated Mutation and Multiple Serine Phosphorylation on Poly(ADP-Ribose) Polymerase 2

**DOI:** 10.1101/2025.04.29.651341

**Authors:** Bridget Hughes, Shubham Chatterjee, Mozhdeh Ghafari, G. Andrés Cisneros

## Abstract

Poly [ADP-ribose] polymerase 2 (PARP2) plays a crucial role in DNA repair. A common single-nucleotide polymorphism (SNP) in the PARP2 gene, rs3093921, has been associated with pancreatic cancer and marginal zone lymphoma (MZL). This SNP results in a missense mutation, D235G, in the PARP2 protein. PARP2 is also reported to undergo post-translational modifications (PTMs), particularly phosphorylation at serine residues 226, 232, and 353. The C–terminal region of PARP2 includes the Trp-Gly-Arg (WGR) and ADP-ribosyl transferase (ART) domains, and the helical subdomain (HD). The latter two, spanning residues 220 to 583, comprise the catalytic region of PARP2. The DNA-induced enzymatic activation of PARP2 is regulated by local destabilization of the HD domain. We used molecular dynamics (MD) simulations to investigate the impact of these three PTMs on both the wild-type (WT) and the mutant (D235G) forms of PARP2. Our simulations suggest that, while neither the cancer–associated mutation nor PTMs on their own significantly alter the overall flexibility of residues within the HD domain, they cause notably greater deviations in the backbone structure of the HD domain compared to the WT. In addition, PTMs in the context of the mutation show reduced interactions between the mutation site and other protein regions, likely due to structural stabilization induced by the PTMs. Importantly, PTMs mitigate the structural disruption caused by the mutation, helping the mutant protein retain a WT-like conformation. Additionally, the HD domain contributes to maintaining PARP2 in its inactive state through its connection with the ART domain. However, the D235G mutation weakens this connection. While PTMs alone do not have a significant impact on this connection, the simultaneous presence of both the mutation and PTMs partially alleviates the disruptive effect of the mutation and leads to partial restoration of the connection between the HD and ART domains.

## Introduction

DNA damage occurs frequently in human cells, with approximately one million damages generated per cell per day [1]. These damages can cause different diseases, such as cancer. To counteract this genomic instability, humans have evolved multiple DNA damage response (DDR) pathways. Poly(ADP-ribose) polymerase 2 (PARP2) is a DNA-binding protein that is involved in multiple DNA repair pathways, including double-strand break (DSB) repair [2, 3, 4], base excision repair (BER) [5, 6], and single-strand break repair (SSBR) [7]. PARP2 helps to repair breaks in DNA by synthesizing poly(ADP-ribose) (PAR) chains using NAD^+^ as a substrate. The highly charged PAR chains promote chromatin relaxation and recruit additional DNA repair proteins to damaged sites. Within the PARP family, PARP1, PARP2, PARP5A, and PARP5B possess the capability to synthesize PAR chains, with PARP1 and PARP2 contributing approximately 80% and 5–15% of total PARP enzymatic activity, respectively.[8, 9, 5, 10, 11, 12, 13, 14, 15, 16, 17, 18, 7, 19, 20]. The N-terminal region of human PARP2 is intrinsically disordered and is not included in the domain overview of PARP2. PARP2 consists of majorly two domains namely, the Trp-Gly-Arg (WGR), and catalytic (CAT) domain, consisting of helical subdomain (HD) and the ADP-ribosyl transferase (ART) subdomain (see Figure 1). The WGR domain of PARP2, which is 33% identical to the same domain in PARP1 [21, 22], plays a critical role in recognizing DNA damage and binding the damaged DNA. The ART domain has ADP-ribosyl transferase activity, featuring a catalytic triad and a binding site for PARP1/2 inhibitors. Its activity is tightly regulated by the HD subdomain; in the absence of DNA, the HD is folded against the ART domain, physically blocking access to the active site and preventing NAD^+^ binding [23, 24, 25, 26]. Like PARP1, PARP2 and PARP3 regulate DNA-induced enzymatic activation by locally destabilizing the HD domain.[24, 27, 28, 29]. Upon DNA binding, the WGR domain initiates structural changes that propagate through the HD domain, causing it to shift away from the ART domain. This conformational rearrangement opens the catalytic site, allowing NAD^+^ to bind and enabling enzymatic activity. This sequence of events forms an allosteric communication pathway between the DNA-binding interface and the catalytic core, ultimately leading to the activation of PARP2 [24, 27, 28, 29, 30, 23, 31].

**Figure 1.**
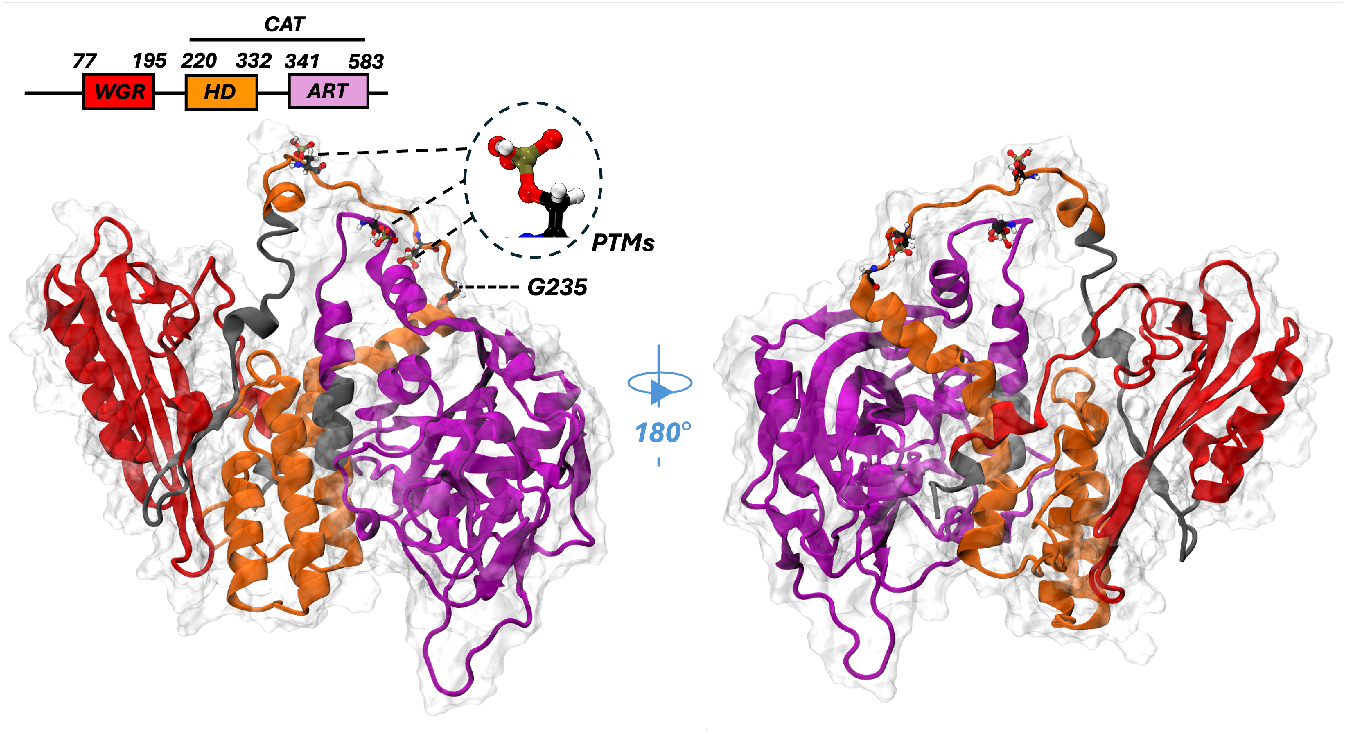
Structural and multidomain organization of PARP2 (PDB ID: 4ZZX). The mutation site and the PTMs are indicated in the figure.

PARP2 has been implicated in various cancers [32, 33, 34, 35, 36]. Previously, we used hypothesis-driven SNP search (HyDn-SNP-S) to identify a common single-nucleotide polymorphism (SNP) in the PARP2 gene, rs3093921 [37, 38], which is strongly associated with pancreatic cancer and marginal zone lymphoma (MZL).[39] This SNP leads to a missense mutation (D235G) in the PARP2 protein. D235 is a conserved residue in the HD domain (Figure 2). Additionally, PARP2 is subject to post-translational modifications (PTMs), namely phosphorylation, at residues S226, S232, and S353.[40]. To our knowledge, these PTMs have not been previously associated with any diseases or cancers. In a previous study [41], phosphorylation of PARP1 has been shown to increase its activity relative to its dephosphorylated state. Understanding how post-translational modifications (PTMs) at positions 226, 232, and 353, as well as the cancer-associated mutation D235G, affect PARP2 could offer insights for PARP2 inhibition.

**Figure 2.**
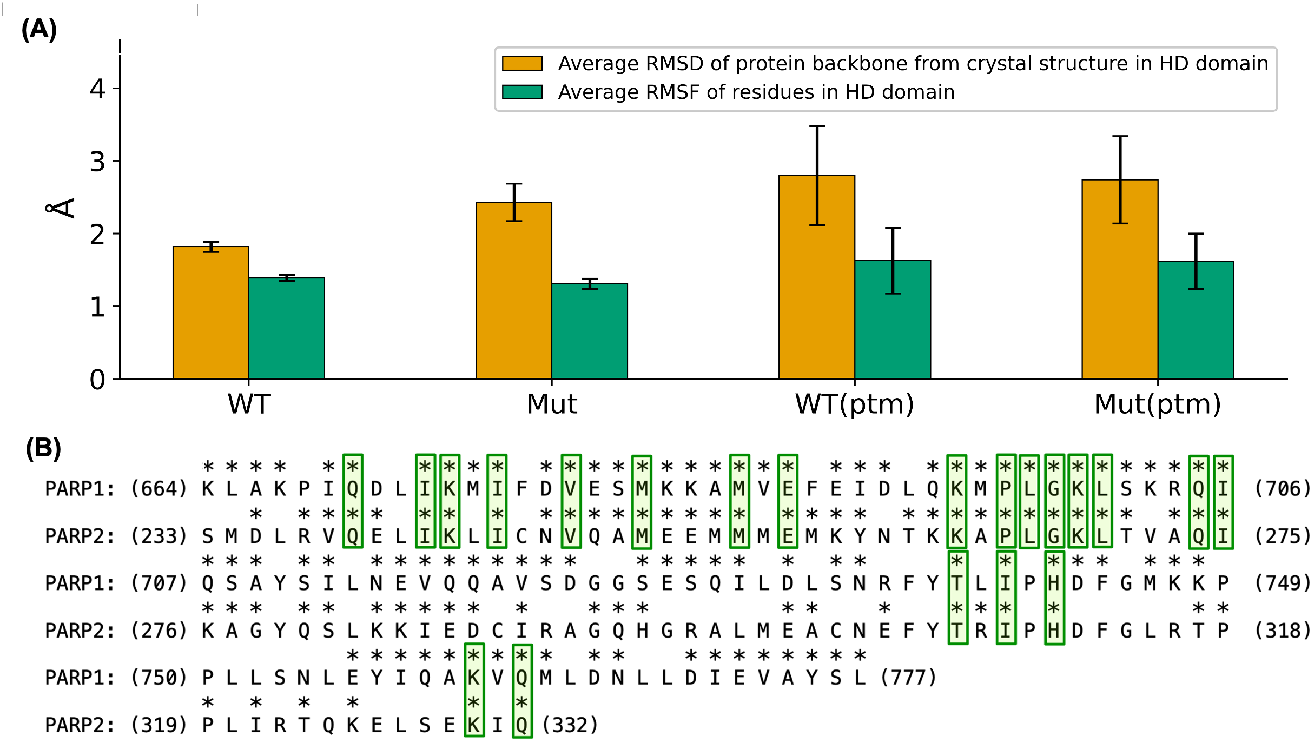
(A) Average time-dependent RMSD (in orange) for the C*α* atoms and average residual RMSF (in green) in the HD domain with respective standard deviations for WT, Mut, WT(ptm), and Mut(ptm) systems. Each calculation used the corresponding initial PDB structure as the reference. (B) Structure-based sequence alignment of the HD domains of PARP1 and PARP2. Asterisks (*) indicate conserved residues, and residues conserved in both PARP1 and PARP2 are boxed in green.

In this study, we investigated the effects of the mutation on PARP2, and the effects of the three PTMs on both wild-type (WT) and mutant (D235G) PARP2 using molecular dynamics (MD) simulations. We refer to the PTM-modified wild-type protein system as WT(ptm) and the PTM-modified mutant system as Mut(ptm). The remainder of the paper is organized as follows: the next section outlines the system setup and details of the MD simulations, followed by a description of the analysis methods. We then present and discuss the results of these analyses, which include time-dependent root-mean-square deviation (RMSD), root-mean-square fluctuation (RMSF), sequence alignment, energy decomposition analysis (EDA), distance analysis, principal component analysis (PCA), dynamic network analysis, and dynamic cross-correlation (DCC), followed by concluding remarks.

## Materials and Methods

### Systems setup for simulations

The native PARP2 crystal structure (PDB ID: 4ZZX) [42] was obtained from the RCSB Protein Data Bank. Missing residues in the structure were modeled using AlphaFold [43, 44, 45, 46, 47, 48]. Specifically, the modeled regions included the WGR domain, parts of the HD domain, and select residues in the ART domain—namely, residues M70 to E231, as well as P549 and D550. Good agreement was observed using UCSF-Chimera [49], between the crystal structure and the AlphaFold2-generated model (see Figure S1). The expected error in the residual position of the modeled PARP2 and the crystal structure remained below 5 Å for the majority of residues (see Figure S1). The rest of the systems—namely, Mut, WT(ptm), and Mut(ptm) were generated from the WT PARP2 structure using UCSF Chimera [49]. Next, the protonation states for ionizable residues were calculated with ProPKA [50], and hydrogens were added based on the protonation states to reflect their ionization at neutral pH. Additionally, atomic clashes were checked and resolved using MolProbity [51]. The LeAP [52] module of AMBER18 [53, 54] was used to neutralize the system with 150 mM KCl and solvate the system with water molecules in a cubic box extending at least 12 Å from the protein surface.

### Simulation details

The ff14SB [55] force field, and the TIP3P [53, 54] water model were used to represent the systems. To relax the system before production, we performed minimization and equilibration steps. The minimization step was performed at 0 K with 5000 steps, where, the initial 1000 minimization steps were performed with steepest descent algorithm, and the remaining steps were preformed with conjugate gradient algorithm. During the minimization oxygen atoms of water were restrained with 100.0 *kcalmol*^*−*1^ Å^*−*2^ restrain. After minimization, the restrain was gradually decreased with short 500 ps MD simulations untill 10.0 *kcalmol*^*−*1^ restrain in steps (100.0, 90.0, 80.0,.., 10.0 *kcalmol*^*−*1^ Å^*−*2^). Next, the systems were gradually heated to 300K with Langevin dynamics[56] with a collision frequency of 2 *ps*^*−*1^ untill no restrains remained. Subsequently, the systems were equilibrated under the NPT ensemble for 50 ns with no restraints, with AMBER18 pmemd.cuda [53, 54] engine. Finally, the production calculations for each system were accomplished in 500 ns of NPT ensemble without restraint in triplicate for 500 ns for each replicate, with AMBER18 pmemd.cuda with an integration time-step of 2 fs. Trajectories were saved every 2 ps. All bonds involving hydrogen were treated with the SHAKE [57, 58] algorithm employed in PMEMD, ensuring all bonds involving hydrogen are constrained. Long-range Coulomb interactions were handled with the smooth particle mesh Ewald (PME)[59, 60] method and long-range van der Waals interactions were approximated using the default isotropic correction [61] in AMBER with a 9 Å cutoff for non-bonded interactions.

### Post-production analysis

We used the production portion of the simulation for further analyses. RMSD, RMSF, PCA, EDA, and DCC analyses were performed for all replicates using AMBER’s CPPTRAJ [62] suite of AMBER. The mathematical formulations behind these methods have been detailed in our previous report [11]. Here, we briefly describe the procedures used.

To calculate the average RMSD of a domain or the entire protein in a given replicate, we firstly measured the RMSD of the alpha carbon atoms (C*α*) in the backbone, using the crystal structure as the reference. These values were then averaged over all frames in the replicate. Finally, to compute the overall average across all replicates, we took the mean of these per-replicate averages. A similar approach was used for RMSF calculations. For each replicate, we calculated the RMSF of each residue, again using the crystal structure as the reference. The residue-wise RMSF values were averaged over all residues in the domain or protein. To obtain the final average across replicates, we averaged these per-replicate RMSF values.

Structure-based sequence alignment of HD domains of PARP1 and PARP2 was carried out with 100 PDBs of the two proteins. An clustal alignment format file (aln) files for the PARPs were generated with BLAST [63, 64, 65, 66, 67, 68, 69, 70] programs, and then these files were used to get the conservation score with ConSurf 2016 [71].

PCA was performed to investigate conformational changes in the protein structure during production runs. PCA is a linear transformation technique applied to atomic coordinates that reduces the system’s dimensionality by identifying the dominant modes of motion. It captures the directions of greatest variance, known as “principal components,” within the internal atomic coordinates of the trajectory. The process begins with constructing a covariance matrix based on these coordinates, followed by eigenvalue decomposition of the matrix to obtain a set of eigenvectors (representing the principal components) and then a diagonal matrix of corresponding eigenvalues.

Energy decomposition analysis (EDA) was performed for the production part of simulations to investigate the nonbonded (NB) intermolecular interactions (Van der Waals and Coulomb) between two fragments with Amber-EDA [72]. We calculated pairwise interaction energies between the mutation site (D235 in the case of WT and WT(ptm), and G235 in the cases of Mut and Mut(ptm) systems) and the rest of the residues. To find the total interaction energy for the mutation site with the HD domain we calculated the sum of interaction energies across all the residues in the HD domain. For the differences in total interaction energy between the mutation site and the HD-domain in a system and the WT system, we calculate the difference in total interaction energy between these systems. To construct the distance matrix, we calculated the average Euclidean distance between the centers of mass of residues across the production for a replica, and then these values were averaged over all replicates. The code for this analysis is available in GitHub [73].

Dynamic cross-correlation (DCC) was used to analyze the correlation in movement between different residues during production. In a fitted structure, the covariance matrix for the position vectors of two residues is calculated. Then, the cross-correlation matrix elements are calculated by normalizing the covariance matrix of these residues, with the product of their standard deviations ensuring values range from –1 (anti-correlated) to +1 (fully correlated). A difference in DCC can be represented as the difference in correlated and anticorrelated motions between two systems. Here, the WT is considered as the reference.

Dynamic network analysis was performed using the DyNetAn Python package [74, 75] to investigate residue–residue interactions across all systems. In this approach, a network is constructed from nodes and edges, where each node represents a residue and is placed at the C*α* atom for amino acids. An edge connects two nodes if any of their heavy atoms remain in contact (≤ 4.5 Å) for at least 75% of the simulation time, provided the residues are not sequential neighbors in the protein chain. The strength of each edge is determined by the weighted correlation between the corresponding residues, reflecting the probability of information transfer along that connection. More highly correlated residues are considered more likely to interact, resulting in more heavily weighted edges.

## Results and Discussion

RMSD values for the residues for the full protein remained around 4 Å for all systems (Figures S2, and S3). Domain-wise RMSD analysis shows that all three domains exhibit similar RMSDs for the WT and Mut systems. However, when PTMs are considered, a significant increase in RMSD in the HD domain is observed. The average RMSF of residues in the HD domain remained comparable in the four systems (Figure 2). Average residual RMSF for each residue in different systems is provided in SI (Figure S4). Residue-wise fluctuations for the HD domain are consistent with the RMSD results, with increased fluctuations mainly observed for residues 220-245 (Figure S4).

As shown in Figure 2, the average RMSD of the protein backbone in the HD domain of PARP2 increases significantly with the mutation and the PTMs. As previously mentioned, PARP2 regulates DNA-induced enzymatic activation by locally destabilizing the HD domain. Given the structural similarity in the HD domains of PARP1 and PARP2 (Figure 2), the enhanced structural deviation with the mutation and the PTMs suggests that these may affect PARP2’s function to repair DNA damage.

Next, we performed EDA for each system to investigate how the mutation site interacts with the rest of the protein (see Figure S5). The differences in interactions between the mutation site and the rest of the residues are shown in Figure 3. In particular, residue R237 exhibits significantly stronger favorable interactions with the mutation site in WT than the Mut and Mut(ptm) systems (Figure 3). In contrast, EDA data for WT(ptm) vs. WT (Figure 3) reveals unfavorable interactions between PTMs and residue D235. However, PTMs do not considerably alter D235’s interactions with other residues.

**Figure 3.**
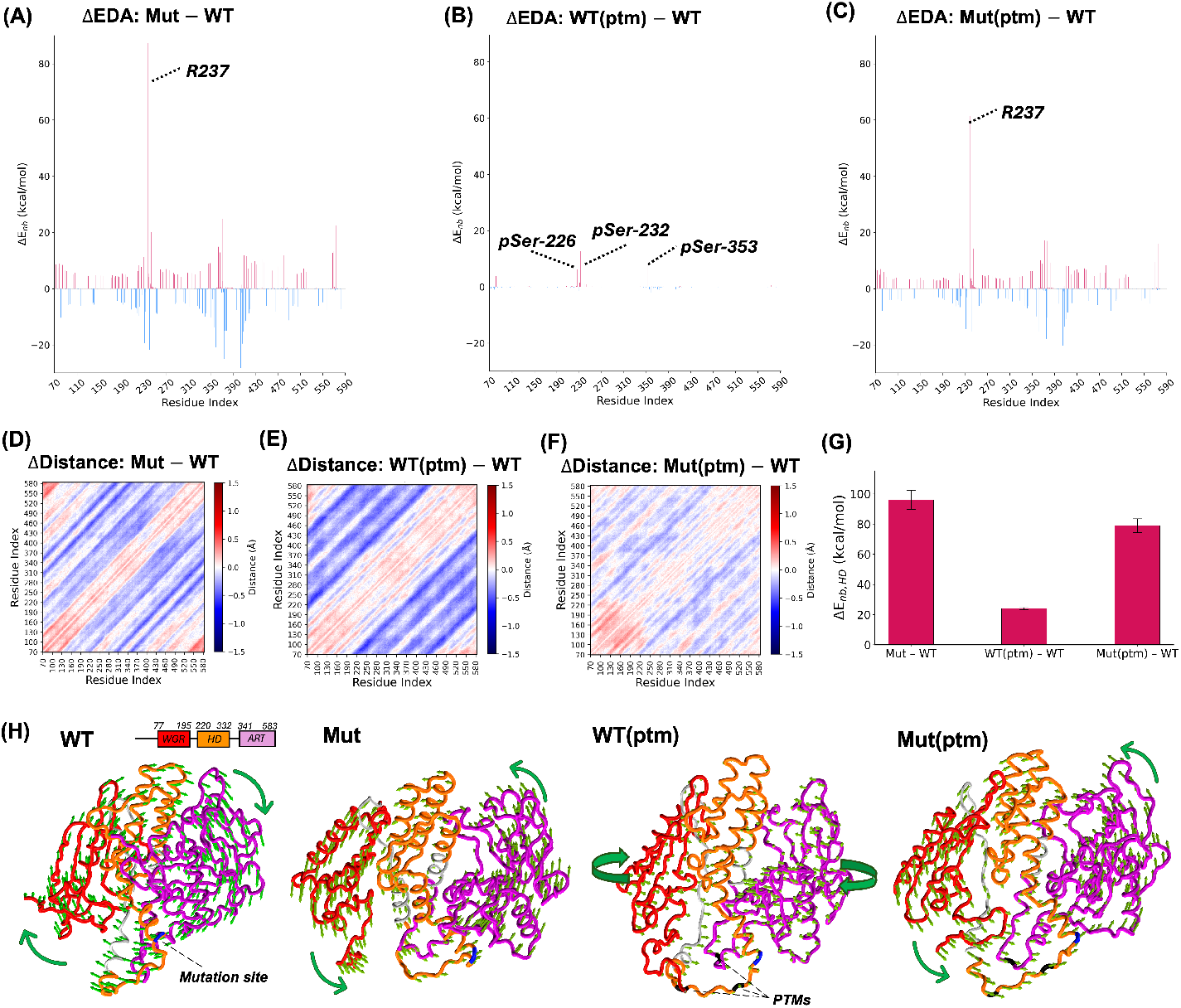
Difference in non-bonded interaction energy between the mutation site and the rest of protein residues between (A) Mut and WT (B) WT(ptm) and WT and (C) Mut(ptm) and WT, with WT as reference in each case. Positive and negative values indicate repulsive and attractive interactions, respectively. Residual distance differences in (D) Mut and WT (E) WT(ptm) and WT and (F) Mut(ptm) and WT, with WT as reference in each system. (G) Sum of the difference in non-bonded interaction energy between the mutation site and the residues in HD domain for the systems, with WT being the reference. Positive values indicate repulsive interactions. (H) Vectors (represented by green arrows) represent the direction of residual motion in the dominant mode of protein motion (mode 1). The mutation site and the PTMs are indicated with blue and black colors, respectively.

The EDA analyses for Mut vs. WT and Mut(ptm) vs. WT (Figure 3) show highly similar patterns, although the interaction magnitudes are reduced in Mut(ptm). This suggests that PTMs in the presence of the mutation dampen the interaction between the mutation site and other residues, potentially due to PTM-induced structural stabilization. To further explore structural effects, we calculated average residue–residue distance matrices over the production trajectories across all replicates (Figures S6 and S7). Figure 3 panels D–G show the differences in these distances for Mut versus WT, WT(ptm) versus WT, and Mut(ptm) versus WT. While both the mutation and PTMs cause noticeable changes in inter-residue distances (Figure 3), the Mut(ptm) system shows minimal structural deviation from WT. This indicates that PTMs may help preserve the WT-like structure by mitigating the structural impact of the mutation.

Given the critical role of the HD domain in DNA repair, we calculated the total interaction energy between the mutation site and the residues within the HD domain. The differences in this interaction energy for Mut vs. WT, WT(ptm) vs. WT, and Mut(ptm) vs. WT are shown in Figure Our results indicate that the mutation introduces significantly unfavorable interactions between the mutation site and the HD domain. Although the presence of PTMs reduces this unfavorable energy, it remains notably high. Interestingly, in the PTM-containing systems, we also observe unfavorable interactions between the residue D235 and the HD domain.

The pronounced deviation in the backbone of the HD domain (Figure 2) and the increased unfavorable interactions with the mutation site (Figure 3) suggest that the HD domain becomes significantly destabilized upon mutation. This observation is consistent with experimental findings that classify this mutation as cancer-associated [39].

The essential dynamics of the protein domains were explored using PCA. Out of the 100 calculated modes, the top 8 were selected for detailed analysis. Mode 1, which represents the dominant motion across all systems, exhibits a rocking-like movement and accounts for over 60% of the total protein motion (Figure S8). In particular, in the WT(ptm) system, the axis of this motion differs from that observed in the WT, suggesting that PTMs may alter the motion-related functions of PARP2. Exploring the functional implications of these differences is beyond the scope of this study. Dynamic cross correlation plots for the WT, Mut, WT (PTM) and Mut (PTM) systems are provided in Figure S9, and the differences in DCC are shown in Figure 4. PARP2 exhibits significant changes in correlated movements within the ART and WGR domains after mutation (Figure 4D). With the introduction of PTMs (Figure 4E), the correlated motions of residues are enhanced within the ART and WGR domains. In the Mut(PTM) system, a further significant enhancement of correlated motions within these domains is observed compared to the WT (Figure 4F). Regarding anticorrelated motions, the mutation significantly alters the interactions between the ART and WGR domains (Figure 4G). The addition of PTMs (Figure 4H) enhances the anticorrelated motions between these domains, and in the Mut(PTM) system, significantly stronger anticorrelated motions between the ART and WGR domains are observed compared to the WT (Figure 4I).

**Figure 4.**
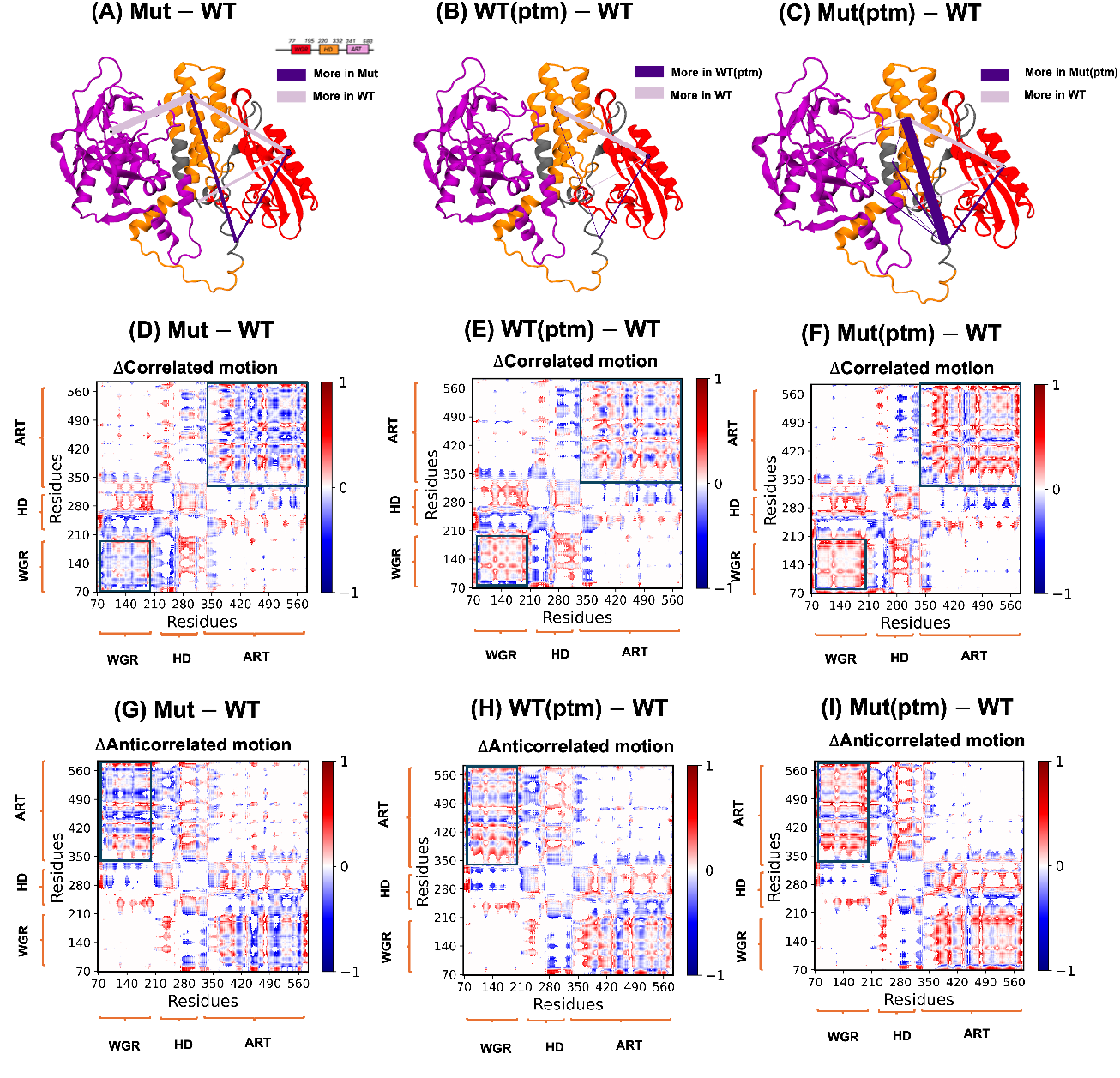
(A-C) Network analysis highlighting the changes in both inter-and intra-domain interactions. The nodes (circles) represent the domains, with their size reflecting the number of interactions occurring within each domain. The edges (lines) connecting the nodes represent interactions between domains, with the line thickness corresponding to the frequency of these inter-domain interactions.(D-F) show the dynamic cross-correlation differences between the Mut, WT(ptm), and Mut(ptm) systems compared to the WT system. (G-I) depict the dynamic cross-anticorrelation differences between the same systems (Mut, WT(ptm), and Mut(ptm)) relative to WT. The results are averaged over three replicates for each system. In the ΔCorrelated motion (Figures 4D–4F) and ΔAnticorrelated motion plots (Figures 4G–4I), negative values indicate stronger correlated and enhanced anticorrelated motions, respectively, in the WT system.

Dynamic network analysis (Figures 4A to 4C) was performed to further investigate the interaction networks within and between domains for each system. The node diagrams presented in this study are organized according to the structural domains to which the residues (nodes) belong. For instance, the circle representing the WGR domain in Figures 4A to 4C indicates the number of intra-domain node–node interactions, reflecting the level of communication within that domain. The thickness of the edges connecting the WGR domain to other domains is proportional to the number of inter-domain interactions. These diagrams were generated by averaging data from three independent replicates for each of the four systems (Mut, Mut(ptm), WT(ptm), and WT). Data for each system are provided in Table S1 and Figure S9 and Table S1. Figures 4A to 4C depict difference plots of the analysis, compared to the WT system.

Network analysis reveals that the cancer-associated mutation significantly reduces the HD–ART node-to-node interactions compared to the WT (Figure 4A), as indicated by the thick lavender-colored edge between the HD and ART domains. The lavender color represents a higher level of inter-domain interactions in WT. Interestingly, PTMs alone do not significantly alter this interaction (Figure 4B), as no edge is observed. However, when both the mutation and PTMs are present together, the disruptive effect of the mutation is partially alleviated, and the HD–ART node-to-node interaction is partially restored, as shown by the thin edge between these domains (Figure 4C). Further, the mutation weakens the HD–WGR node-to-node interactions compared to WT, as reflected by the thick ligher-violet edge between the HD and WGR domains (Figure 4A). The presence of PTMs at the specified residues contribute to the reduction of this interaction, as seen in Figure 4B. Interestingly, when both the mutation and PTMs are present together, the HD–WGR interaction remains significantly weakened. Instead, a stronger interaction between the HD domain and the linker region (residues 127–151, shown in grey in Figure 4A–C) emerges, as indicated by the thick purple edge connecting the linker and HD domain.(Figure 4C).

## Conclusion

Understanding the effects of a cancer-associated mutation and PTMs in PARP2 could aid therapeutic development. We observed notable differences in the structure, dynamics, and interactions of PARP2 associated with the mutation, both in the presence and absence of PTMs, analyzed on a case-by-case basis. Our study shows that while the average RMSD and RMSF in the HD domain remains comparable for the WT, Mut, WT(PTM), and Mut(PTM) systems, the RMSD of the protein backbone in the HD domain and the RMSF are considerably higher in the mutated and PTM-modified systems compared to the WT. Residue–wise interaction energy analysis (EDA) suggests that PTMs, in the presence of the mutation, may dampen interactions between the mutation site and the rest of the protein due to potentially PTM-induced structural stabilization. Interactions between the mutation site and the HD domain show that the mutation introduces significantly unfavorable interactions. This is consistent with a residual distance analysis, which suggests that PTMs may help maintain a WT-like structure by mitigating the mutation’s structural impact. Additionally, network analysis showed that the mutation weakens the connection between the HD and ART domains, which are essential for keeping the enzyme inactive. While PTMs alone do not have a significant effect on this connection, when both the mutation and PTMs are present simultaneously the connection is restored, and the disruptive impact of the mutation on this connection is reduced. DCC results indicate that the mutation affects the correlated and anti-correlated motions of residues within the ART and WGR domains, whereas the PTMs primarily influence the correlated and anti-correlated motions between the ART and WGR domains. Moreover, the combination of the mutation and PTMs significantly amplifies the effects observed with PTMs alone. Lastly, the PCA indicates that the mutation does not significantly affect the dominant motions of PARP2, but PTMs may alter its motion-related functional dynamics.

## Author Contributions

BH and GAC designed the project. BH carried out all simulations, analyzed the data, and prepared the initial draft of the article. SC and MG analyzed and interpreted the data and co-wrote the manuscript. SC and MG contributed equally to this work. All authors contributed to the discussion and manuscript editing. The authors declare no competing interests.

## Acknowledgments

This study was funded by NIH Grant No. R35GM151951. Computational time for this project was provided by the NSF Grant No. OAC-2117247, NSF ACCESS Project No. BIO240023, and the University of Texas at Dallas’ Cyberinfrastructure and Research Services, Ganymede, and Titan HPC clusters.

## Supplementary Material

The supplementary data is available in SI-parp2.pdf, and electronic supplementary data can be found at Zenodo repository https://doi.org/10.5281/zenodo.15305015, DOI 10.5281/zenodo.15305015.

## Notes

https://doi.org/10.5281/zenodo.15305015

### Competing Interest Statement

The authors have declared no competing interest.

