## Supplementary material for "Effects of a Cancer-Associated Mutation and Multiple Serine Phosphorylation on Poly(ADP-Ribose) Polymerase 2": SI

#### 1. Alignment between modeled and crystal structure

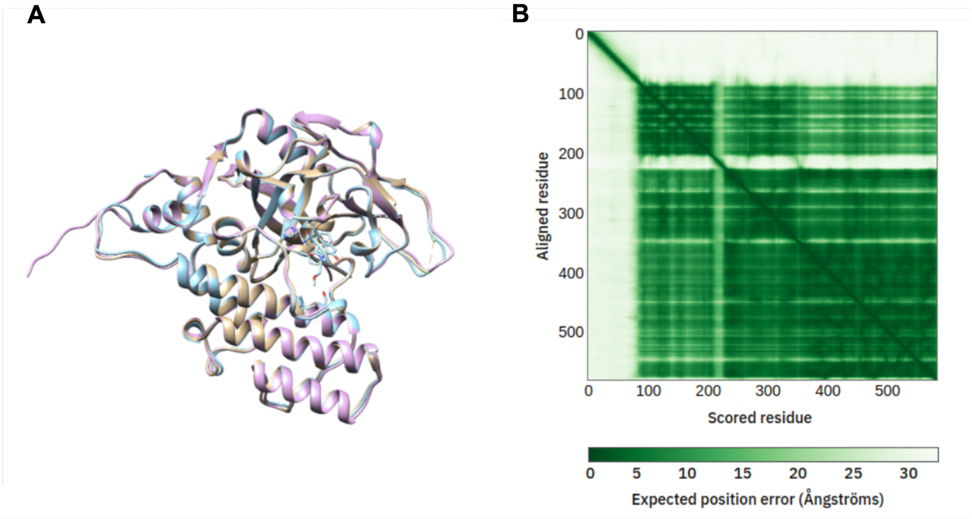


**Figure S1. (A)** Structural overlap/ alignment between the modeled PARP2 structure and PDB ID: 4ZZX (**B**) Expected positional error between the modeled PARP2 structure and PDB ID: 4ZZX

#### 2. RMSD

##### 2.1. All the domains

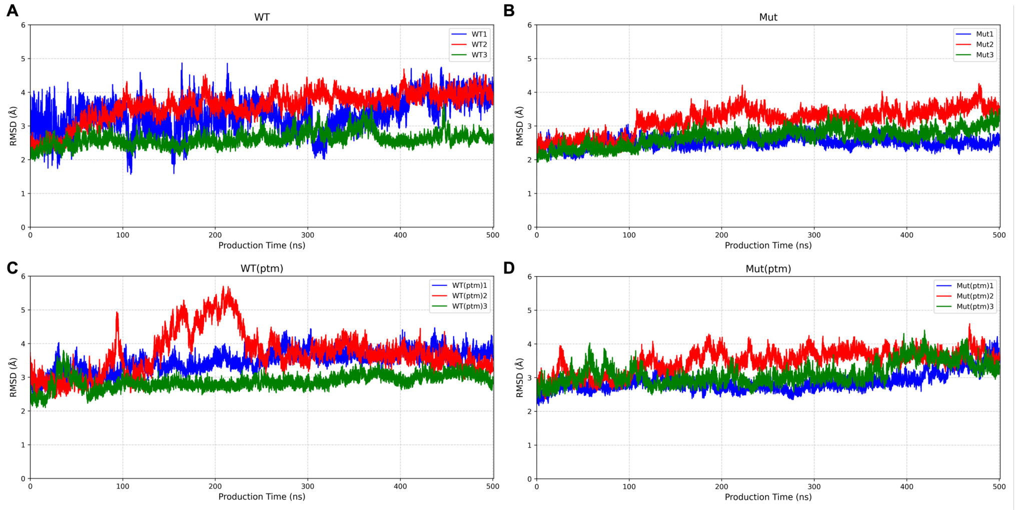


**Figure S2.** RMSD of protein backbone in **(A)** WT **(B)** Mut **(C)** WT(ptm) and **(D)** Mut(ptm) systems, with respect to the crystal structure.

#### 2.2. HD domain

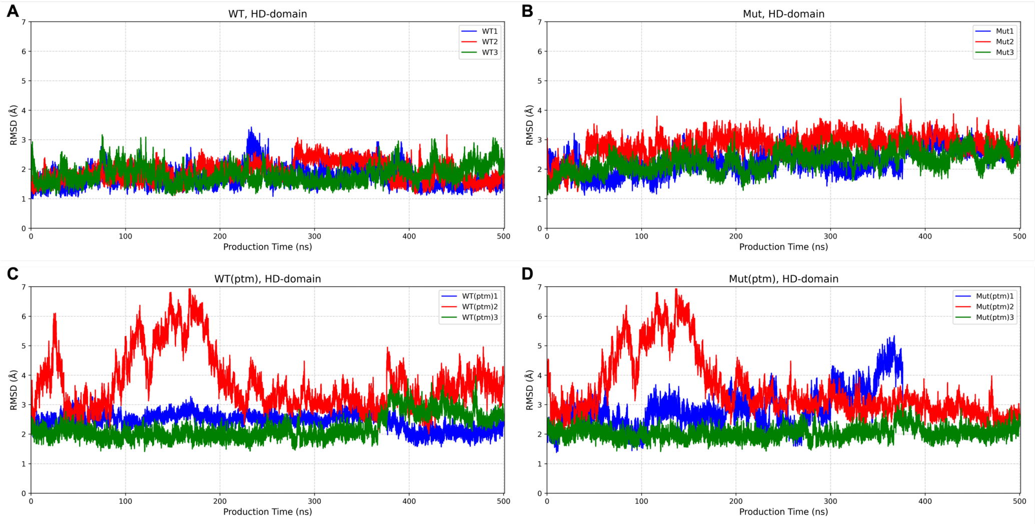


**Figure S3.** RMSD of protein backbone in the HD domain of **(A)** WT **(B)** Mut **(C)** WT(ptm) and **(D)** Mut(ptm) systems, with respect to the crystal structure.

#### 3. RMSF for HD domain

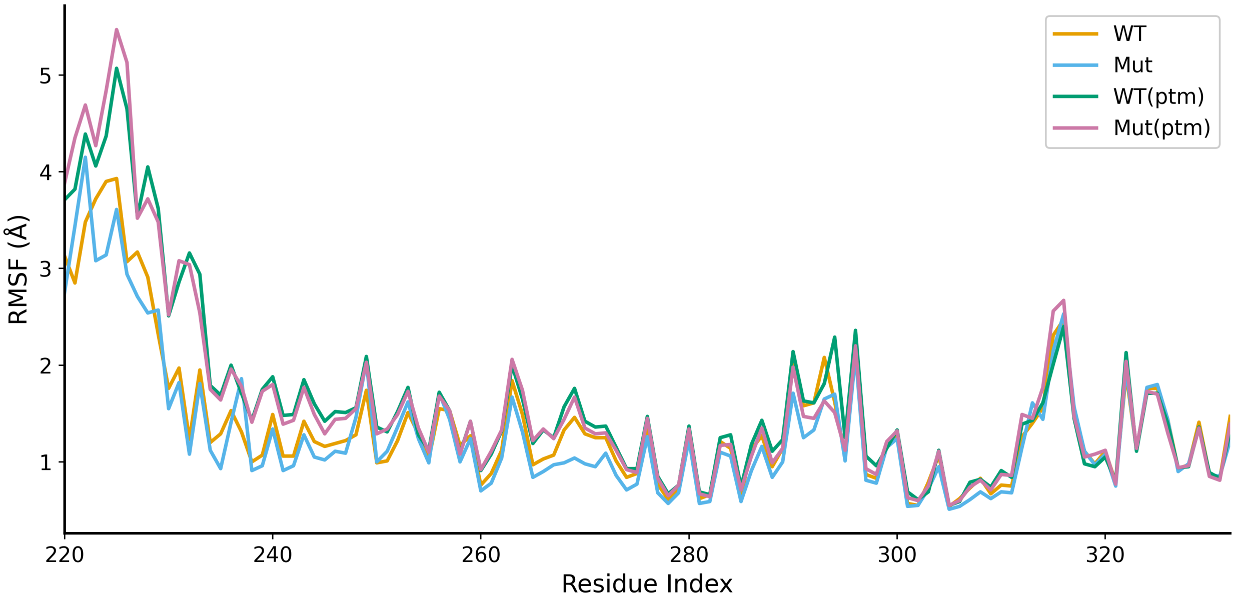


**Figure S4.** Average RMSF of protein residues across all the replicates in the HD domain of **(A)** WT **(B)** Mut **(C)** WT(ptm) and **(D)** Mut(ptm) systems, with respect to the crystal structure.

#### 4. EDA

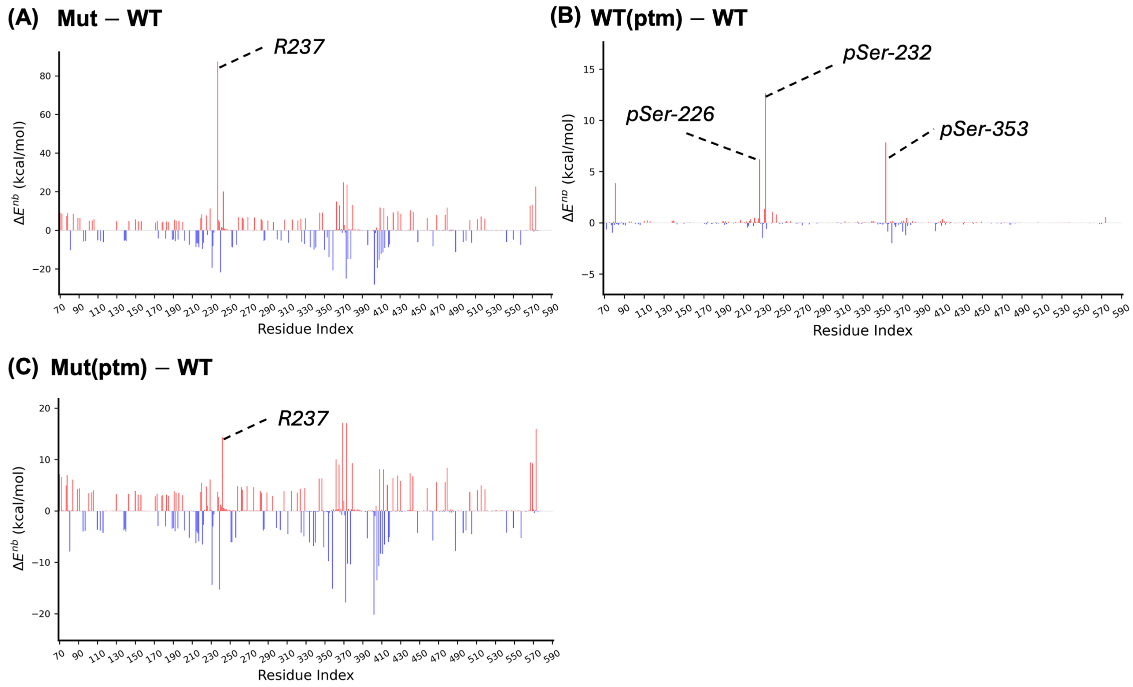


**Figure S5.** Change in average non-covalent interactions between the mutation site (D235 in the the case of WT, and G235 in the case of Mut) with rest of the protein residues.

### 5. Distance matrix and corresponding standard deviation matrix

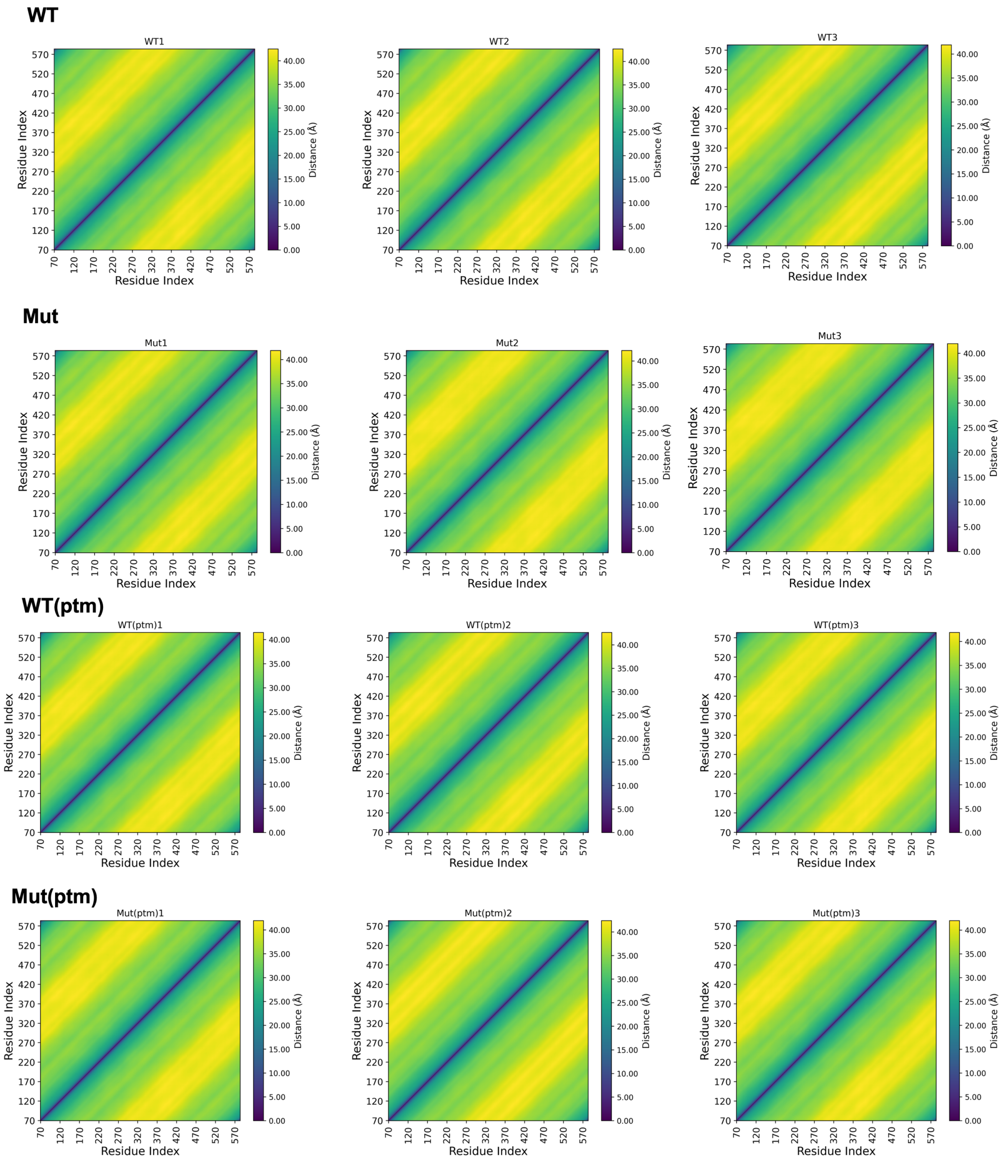


**Figure S6.** Residual distances in replicates of each system.

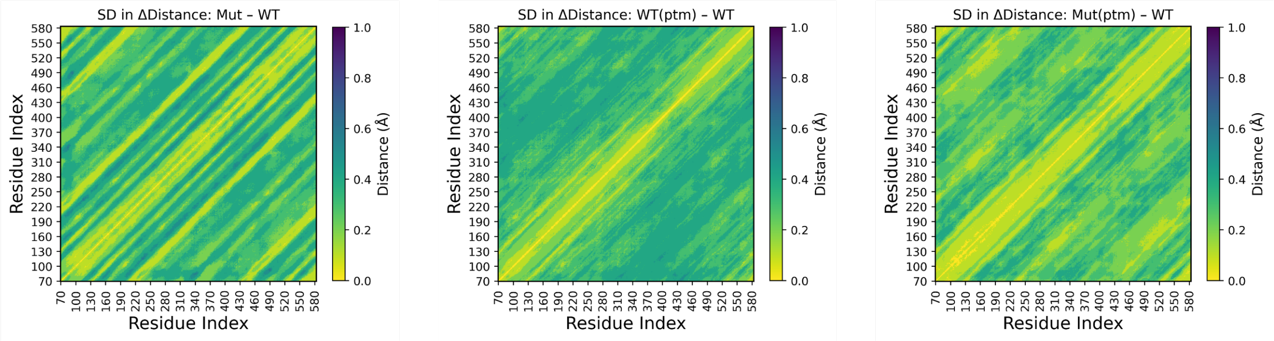


**Figure S7.** Standard deviation in distance in residual distances between residues in Mut, WT(ptm) and Mut(ptm) with WT as reference.

#### 6. PCA

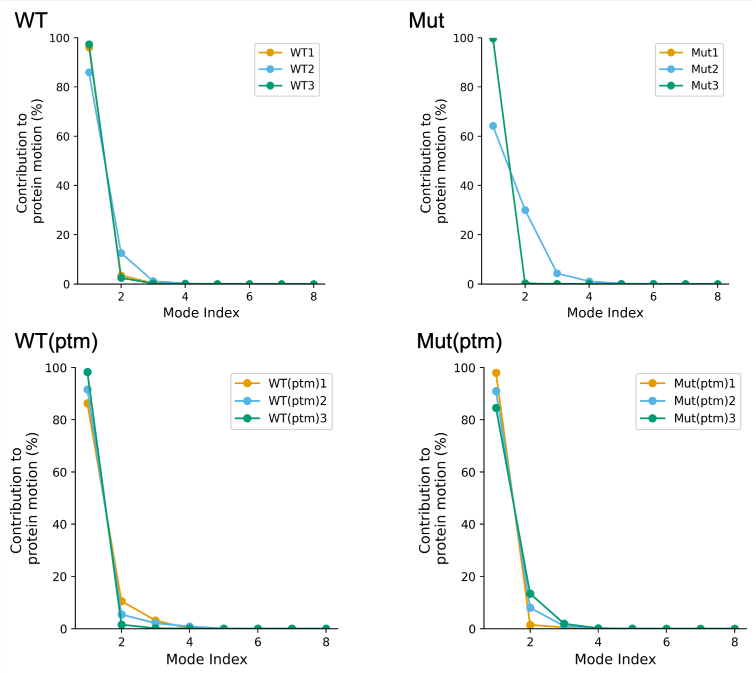


**Figure S8.** Change in average non-covalent interactions between the mutation site (D235 in the the case of WT, and G235 in the case of Mut) with rest of the protein residues.

#### 7. Dynamic Network Analysis

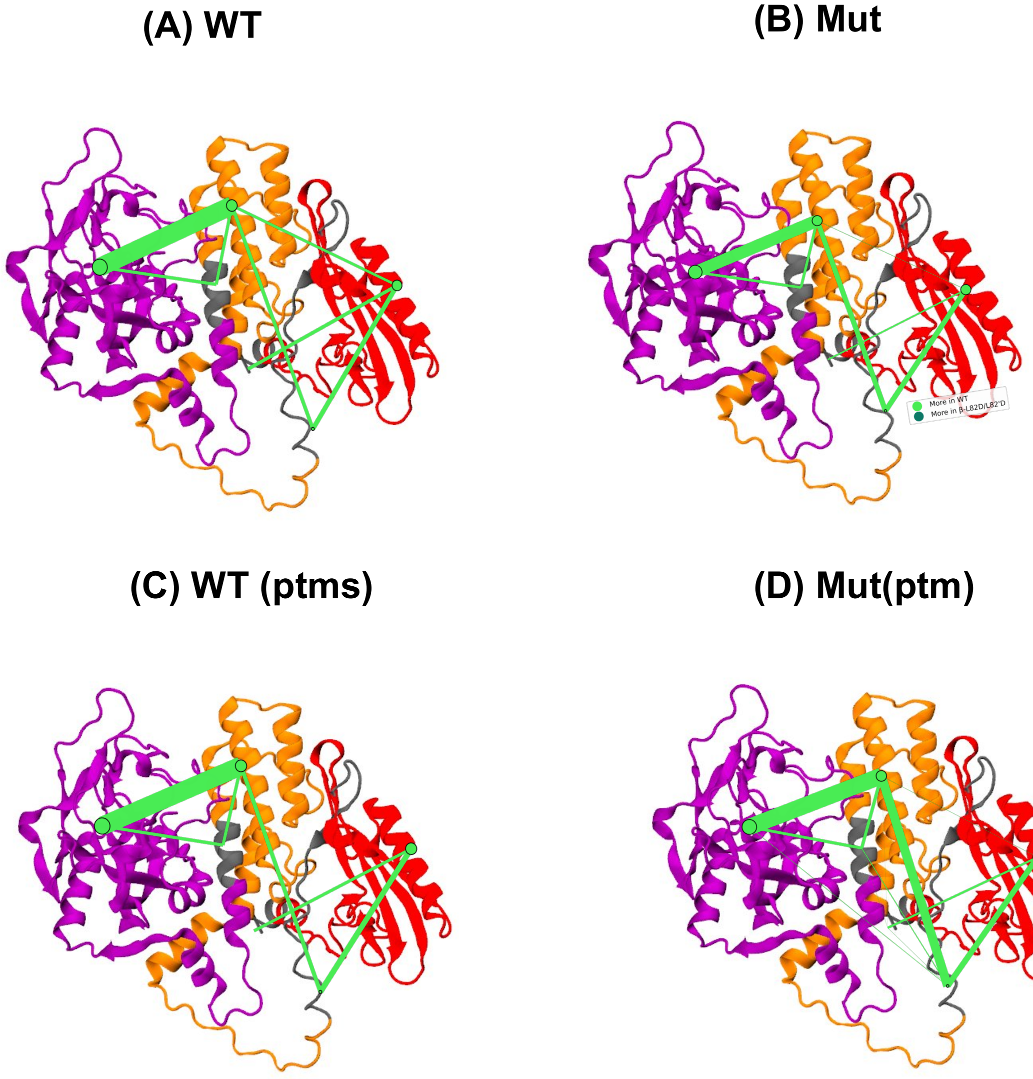


**Figure S9**. The network analysis illustrates changes in both inter- and intra-domain interactions. Nodes (circles) represent individual domains, with their size indicating the number of intra-domain interactions. Edges (lines) between nodes represent inter-domain interactions, with line thickness reflecting their frequency.

| Interaction Type | WT | WT(ptm) | Mut | Mut(ptm) |
|---|---|---|---|---|
| N-terminal | 0 | 0 | 0 | 0 |
| WGR | 678 | 801 | 869 | 638 |
| Linker1 | 60 | 56 | 53 | 51 |
| HD | 802 | 817 | 850 | 796 |
| Linker2 | 0 | 0 | 0 | 0 |
| ART | 1567 | 1557 | 1548 | 1541 |
| N-terminal and WGR | 44 | 38 | 25 | 30 |
| N-terminal and Linker1 | 0 | 0 | 0 | 3 |
| N-terminal and HD | 0 | 0 | 0 | 1 |
| N-terminal and Linker2 | 0 | 0 | 0 | 2 |
| N-terminal and ART | 0 | 0 | 0 | 0 |
| WGR and Linker1 | 59 | 69 | 75 | 77 |
| WGR and HD | 35 | 0 | 6 | 4 |
| WGR and Linker2 | 0 | 0 | 0 | 0 |
| WGR and ART | 0 | 0 | 0 | 0 |
| Linker1 and HD | 47 | 51 | 73 | 120 |
| Linker1 and Linker2 | 0 | 0 | 0 | 12 |
| Linker1 and ART | 0 | 0 | 0 | 7 |
| HD and Linker2 | 40 | 40 | 40 | 43 |
| HD and ART | 220 | 226 | 176 | 218 |
| Linker2 and ART | 40 | 40 | 40 | 40 |

**Table S1**. Breakdown of intra- and intra-molecular interactions in the WT, WT(ptm), Mut and Mut(ptm) systems. This data was extracted from the network analysis and represents averages from tree replicates for each system.

#### 8. DCC

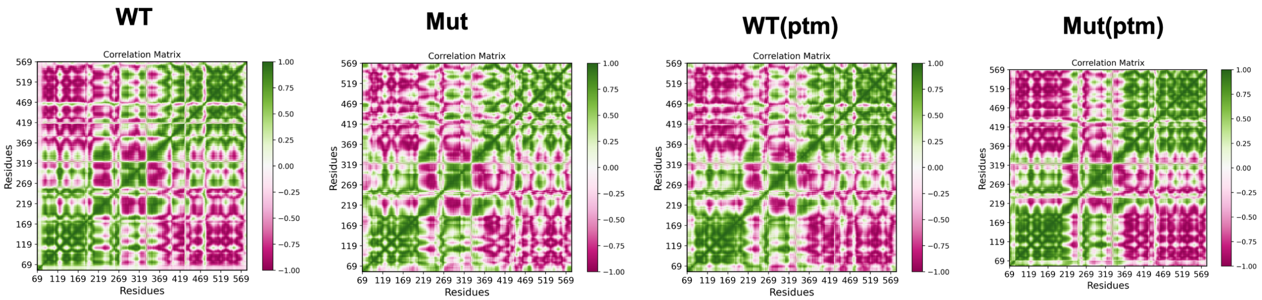


**Figure S10.** DCC plots for the WT, Mut, WT(ptm) and Mut(ptm) systems.
